# Normative Modeling of Static and Dynamic Functional Connectivity

**DOI:** 10.64898/2026.04.03.716292

**Authors:** Nina Baldy, Paul Triebkorn, Spase Petkoski, Meysam Hashemi, Viktor Jirsa

## Abstract

The transition toward individual-level modeling in functional neuroimaging is severely bottlenecked by methodological heterogeneity, which confounds subject-specific effects with study-dependent variance. Normative modeling leverages a multi-level statistical model to parameterize and account for such variability. We examine whether this framework can harmonize fragmented legacy datasets from seven large cohorts (*N* = 4705), bypassing the need for intensive reprocessing under an arbitrary unified pipeline. We evaluate the construction of a lifespan normative chart by harmonizing open-access functional MRI data extracted with highly heterogeneous processing pipelines, incorporating study-level random effects within a generalized additive model. From this well-calibrated model we drive normative trajectories of both static and dynamic functional connectivity and quantify their invariance to the choice of metric used to compute the interactions between regions. We observe a key decoupling between static and dynamic connectivity. While static connectivity exhibits a monotonic age-related decline, dynamic connectivity follows a more complex trajectory: decreasing during pediatric stabilization, peaking in mid-adulthood metastability, and ultimately declining into senescent rigidity. Overall, this framework establishes a scalable statistical blueprint for modeling functional brain organization to circumvent massive data homogenization, while preserving sensitivity to both between-subject variability and within-subject temporal dynamics.

## 1. Introduction

The paradigm of neuroimaging research is undergoing a fundamental shift from group-level case-control studies toward normative modeling, which quantifies individual deviations relative to a population distribution (Marquand et al., 2016; Rutherford et al., 2023). This methodological pivot stems from a growing recognition of the substantial biological heterogeneity that exists within clinical populations (Segal et al., 2025). Historically, research has relied heavily on case-control designs, which implicitly assume that individuals sharing a common diagnostic label also share a uniform neurobiological signature. However, mounting evidence suggests that the manifestation of psychiatric and neurological disorders is highly patient-specific (Wolfers et al., 2018; Zabihi et al., 2020). In many cases, the neurobiological distributions of patients and healthy controls overlap to such a degree that group-level averages fail to capture the underlying pathology, challenging the validity of the “average patient” assumption (Marquand et al., 2016, 2019).

To address these limitations, the field has increasingly adopted normative modeling frameworks to map individual deviations from a population reference, analogous to pediatric growth charts for height and weight (Borghi et al., 2006). This approach has proven particularly robust in structural neuroimaging, where the aggregation of over 120000 scans has established stable lifespan trajectories for cortical thickness, surface area, subcortical volume (Bethlehem et al., 2022), and asymmetry (Dorfschmidt et al., 2025), providing a benchmark for quantifying heterogeneity in disorders ranging from schizophrenia to Alzheimer’s disease (Wolfers et al., 2018; Loreto et al., 2024; Shafiei et al., 2025; ENIGMA Clinical High Risk for Psychosis Working Group et al., 2024; Haukvik et al., 2025). The unprecedented scale of these structural charts relied on hierarchical statistical harmonization, where random effects were leveraged to account for methodological and technical heterogeneity (Bayer et al., 2022).

Indeed, the challenge of methodological heterogeneity arising from even minor disparate data acquisition protocols and analytic pipelines poses a significant challenge in the generalization of normative models (Botvinik-Nezer et al., 2020; Li et al., 2024). Downstream decisions about motion correction, physiological denoising, and particularly the inclusion or exclusion of global signal regression are known to dramatically alter the distributional properties of functional connectivity matrices (Murphy and Fox, 2017; Power et al., 2014). Worse, even upstream rigorously standardized initiatives, such as the Human Connectome Project (HCP) have shown systematic discrepancies across cohorts arising from modest protocol deviations (Chan et al., 2026). While retrospective data harmonization techniques like ComBat (Johnson et al., 2007; Fortin et al., 2017, 2018) are widely used to explicitly regress out site-specific batch effects, these approaches frequently over-correct, inadvertently stripping away intrinsic biological variability alongside measurement-related noise. In contrast, hierarchical Bayesian modeling incorporates site effects directly into the generative architecture. As demonstrated by Bayer et al. (2022), this generative modeling approach vastly outperforms explicit harmonization techniques in both predictive power and the preservation of biological variance; notably, their evaluation revealed that ComBat discarded over 90% of the original variance, severely degrading downstream predictions.

Building on the structural foundation by Bethlehem et al. (2022), the frontier of normative modeling has recently expanded to the functional connectome. A landmark study by Sun et al. (2025) established the first comprehensive growth charts for the human functional connectome, aggregating data from over 33000 individuals across 132 sites. Their work revealed that the global mean of functional connectivity follows a non-linear trajectory, peaking in the late fourth decade of life, while connectivity variance peaks earlier, in the late third decade, highlighting a temporal dissociation between central tendency and dispersion. This study employed a rigorous two-step harmonization strategy. First, pipeline-related variance was mitigated by reprocessing all raw data through a uniform minimal pipeline (alongside developing a lifespan-wide suite of system-level atlases). Second, a random-effects framework was applied to account for residual site-and scanner-related variance. Other initiatives of unified large-scale data sharing include the Reproducible Brain Charts (Shafiei et al., 2025), which assembled data from five major developmental cohorts and employed bifactor modeling to derive generalizable developmental dimensions from both structural and functional perspectives. This approach handled variability by using a unified and reproducible pipeline, followed by CovBat (Chen et al. (2022); a covariance correcting modification of ComBat) harmonization combined with generalized additive models in order to address the residual site-specific batch effects without discarding relevant variability.

While the combination of uniform reprocessing and statistical harmonization applied in previous studies (Sun et al., 2025; Shafiei et al., 2025) can yield high-fidelity references, the prerequisite of monolithic preprocessing imposes an immense computational and resource burden, effectively limiting the generation of functional normative models to large-scale consortia. Furthermore, because these gold-standard initiatives enforce a single preprocessing pipeline prior to model training, the resulting normative references are explicitly conditioned on that specific pipeline. Consequently, a disconnect remains between the construction of these models and their practical deployment: for a secondary analyst or clinician to validly compute a patient’s deviation score, access to raw data and sufficient resources is required to process the subject using the identical preprocessing pipeline and parcellation scheme as the reference cohort. This effectively implies the need for a globally standardized pipeline, a consensus that the functional neuroimaging community has not yet reached.

Further downstream, arises the question of generalization to legacy datasets. These datasets are often fixed in disparate spatial configurations, extending to the very definition of the brain’s parcellation and anatomical organization. The literature is currently fragmented between anatomically defined atlases, such as the Automated Anatomical Labeling (AAL) map (Tzourio-Mazoyer et al., 2002) and functionally derived parcellations, such as the Schaefer atlas (Schaefer et al., 2018). These schemes reflect fundamentally different methodological assumptions. Structural parcellations rely on sulcal and anatomical landmarks which may not correspond to functional boundaries, whereas functional parcellations are defined by signal coherence but lack anatomical rigidity. Rather than enforcing an artificial equivalence between discordant parcellations, we ask is the global aging trajectory of the functional connectome invariant to the methodological lens through which it is observed?

Beyond spatial definitions, the mathematical formulation used to quantify functional connectivity is an additional layer of methodological heterogeneity. While Pearson correlation remains the dominant standard in the field, alternative metrics, ranging from nonlinear information-theoretic measures such as mutual information to spectral coherence and partial correlations, are also employed to capture distinct facets of brain dynamics (Liu et al., 2025). Consequently, it is worth investigating whether the macroscopic trajectory of the aging connectome, and its statistical harmonization, remains stable across these alternative metrics. Crucially, while the normative reference of the functional connectome established by Sun et al. (2025) represents a monumental step forward, it is inherently limited to a specific representation of static functional connectivity–namely, correlation computed over a time-averaged snapshot that obscures the non-stationary nature of the brain. The human connectome is highly metastable, continuously assembling and disassembling functional hubs to support shifting cognitive states (Hansen et al., 2015; Kong et al., 2021; Lavanga et al., 2023). Functional Connectivity Dynamics (FCD) captures this temporal evolution, operationalizing network fluidity as the variance of time-resolved correlation matrices computed across sliding windows (Hutchison et al., 2013; Lurie et al., 2020). Given that dynamic reconfiguration underpins adaptive brain function (Lurie et al., 2020; Fornito et al., 2015; Shine et al., 2016; Zalesky et al., 2014), the developmental and aging trajectory of the connectome’s temporal organization remains a critical and largely unmapped frontier.

This study explores a computationally viable alternative to the centralized reprocessing of raw neuroimaging data. We investigate whether a multi-level probabilistic model can effectively aggregate heterogeneous data sources into a unified normative framework without the computational burden of full re-processing. We leverage a diverse collection of seven datasets (*N* = 4705), including ABIDE (Di Martino et al., 2014), ADHD-200 (HD-200 Consortium, 2012), ADNI (Jack et al., 2008), CAMCAN (Shafto et al., 2014), HCP Young Adult (Van Essen et al., 2013) and HCP Aging (HCA) (Bookheimer et al., 2019), and 1000BRAINS (Caspers et al., 2014), that span the human lifespan but are fragmented across different parcellations (AAL, Schaefer100, and ICA-based) and processing strategies. By explicitly modeling the preprocessing pipeline or choice of atlas as random effects within a hierarchical Bayesian framework, we aim to estimate the underlying biological signal of functional maturation and fluidity that persists across these technical boundaries. This study details the construction of multi-atlas normative charts for both the static connectome and its temporal dynamics, quantifies the sensitivity of developmental trajectories to metric selection (e.g., correlation versus coherence and precision), and critically evaluates the limits of harmonization when ground truth data representations diverge.

## 2. Materials and Methods

### 2.1. Dataset Aggregation

To construct a lifespan normative model of the functional connectome, we aggregated already processed resting-state functional MRI (rs-fMRI) data from seven large-scale, open-access repositories, spanning an age range from childhood to late senescence. The combined cohort included data from (see Table 1 and Figure 1):

- HCP Young Adult (Human Connectome Project, Van Essen et al. (2013));
- HCP Aging & AABC (Aging Adult Brain Connectome, Bookheimer et al. (2019));
- 1000BRAINS (conducted at the Research Centre Julich, Caspers et al. (2014));
- CAMCAN (Cambridge Centre for Ageing and Neuroscience, Shafto et al. (2014));
- ABIDE I & II (Autism Brain Imaging Data Exchange, Di Martino et al. (2014); preprocessed from the Neuro Bureau Preprocessing Initiative, Craddock et al. (2013));
- ADHD-200 (Attention Deficit Hyperactivity Disorder, HD-200 Consortium (2012); preprocessed from the Neuro Bureau Preprocessing Initiative, Craddock et al. (2013));
- ADNI (Alzheimer’s Disease Neuroimaging Initiative, Jack et al. (2008)).

**Figure 1.**
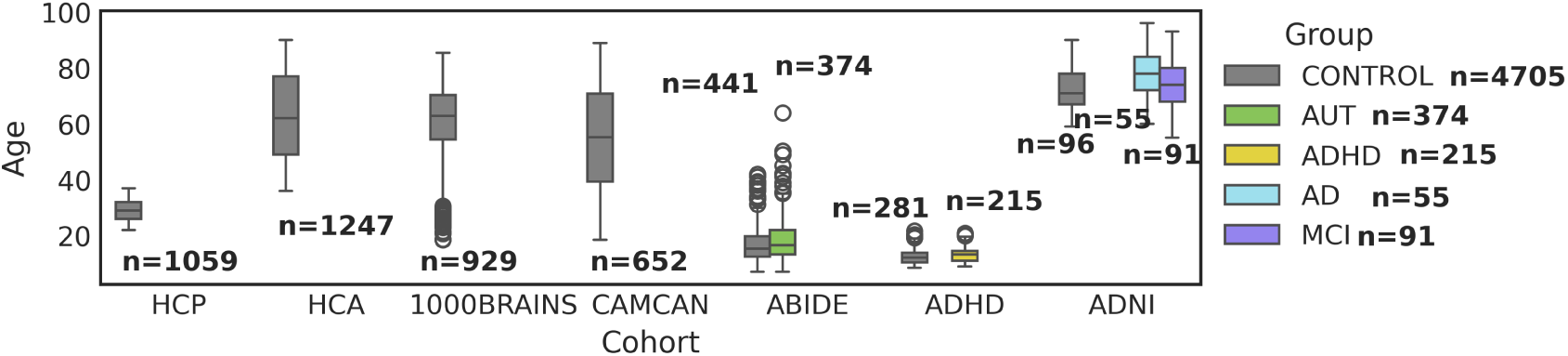
Overview of the aggregated dataset used in this study, from seven large-scale open-access repositories, covering the human lifespan from childhood to late senescence.

**Table 1.** Summary of of demographic, diagnostic, and parcellation characteristics of the aggregated neuroimaging datasets used in this study.

| Dataset | N control | N case | Age median (range) | Sex | Parcellation |
| --- | --- | --- | --- | --- | --- |
| HCP Young Adult | 1059 | 0 | 29 (22-35) | 54.4% F | Schaefer100 & AAL |
| HCP Aging (HCA) - AABC | 1247 | 0 | 61 (36-90) | 57% F | Schaefer100 & AAL |
| 1000BRAINS | 929 | 0 | 62.8 (18-85) | 43% F | Schaefer100 |
| CAMCAN | 652 | 0 | 55.2 (18-89) | 50.6% F | AAL |
| ABIDE | 441 | 374 AUTISM | 14.8 (6-64) | 20.8% F | AAL |
| ADHD | 281 | 215 ADHD | 10.8 (7-22) | 35% F | AAL |
| ADNI | 96 | 55 AD, 91 MCI | 74 (55-96) | NA | Schaefer100 |

Collectively, these datasets represent a large aggregation of the widely used open-access rs-fMRI data, characterized by varying acquisition protocols and sites, scanner manufacturers, demographics, preprocessing pipelines, and parcellation atlases. Control data (*N* = 7011, with *N* = 4705 unique subject IDs, the *N* = 2306 HCP/HCA subjects are represented in two atlases) was partitioned into healthy train (*N* = 4907) and test sets (*N* = 2104) in proportions 70% and 30%, stratified over dataset and atlas. Case data was only used as a test case set (*N* = 770).

### 2.2. Data Processing

A central aim of this study was to evaluate the robustness of normative trajectories to methodological variation. Accordingly, rather than reprocessing all raw data through a single monolithic pipeline, we leveraged the precomputed time series available from each consortium, resulting in a dataset stratified by distinct methodological factors. Specifically, the data were available in either a structural parcellation: the Automated Anatomical Labeling (AAL) atlas (90 or 116 regions), defined by sulcal landmarks (Tzourio-Mazoyer et al., 2002), or a functional parcellation: the Schaefer-100 atlas, defined by local-global functional clustering (Schaefer et al., 2018).

To statistically estimate the variance attributable to atlas choice, the HCP consortium was treated as a reference bridge. We obtained time series from the HCP Young Adult and HCP Aging datasets, extracted using both the AAL and Schaefer atlases. Because the site and scanner are held constant while the parcellation varies, this overlap allows the hierarchical model (see subsection 2.4) to disentangle age-related fixed effects from atlas-specific random effects.

### 2.3. Features of Functional Connectivity

For each subject, we computed metrics of static Functional Connectivity (FC; Friston (1994); Honey et al. (2009); Greicius et al. (2003)) using different metrics, reflecting the statistical dependence between brain region, and Functional Connectivity Dynamics (FCD; Zalesky et al. (2014); Hansen et al. (2015); Lurie et al. (2020)) using sliding-window approach, reflecting time-varying fluctuations in these inter-regional interactions.

#### 2.3.1. Static Connectivity Metrics

Recent benchmarking efforts highlight that no single metric optimally captures all dimensions of functional network topology (Liu et al., 2025). We aimed to explicitly quantify how the choice of connectivity metric influences the estimated lifespan trajectory, and for doing so, we extracted FC using several distinct estimators:

- Pearson correlation: The standard measure of linear dependence.
- Spearman’s rank correlation: A rank-based measure robust to non-Gaussian signal distributions.
- Mutual information: An information-theoretic measure quantifying the statistical dependence between two signals.
- Magnitude-squared coherence: A frequency-specific measure of coupling (averaged 0.01–0.1 Hz).
- Precision (partial correlation): Estimated via the inverse covariance matrix.

An illustration of FC matrices obtained with these metrics is given in Figure 2. For each metric, the upper triangular elements of the matrix were averaged to obtain a single scalar representing the global connectivity strength for each subject.

**Figure 2.**
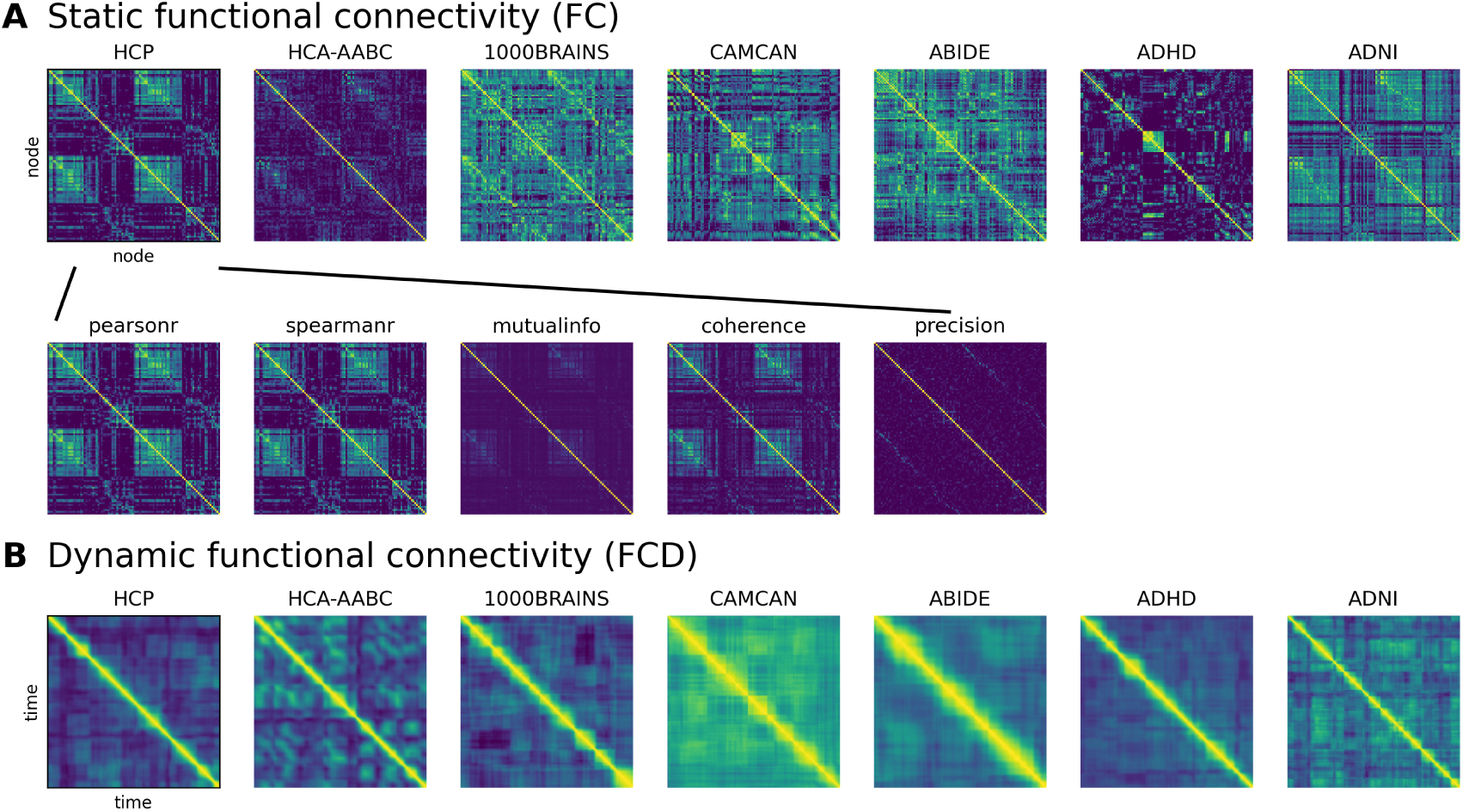
Visualization of functional connectivity and functional connectivity dynamics. (**A**) Shown are static functional connectivity (FC) across cohorts, computed as pearson correlation between nodes (regions) of the parcelated brain. Other metric-specific FC matrices are shown for a representative individual in the HCP dataset. The metrics span standard statistical measures (Pearson and Spearman’s rank correlations, precision matrix), information-theoretic measures (mutual information), and spectral measures (magnitude-squared coherence). (**B**) Dynamic functional connectivity (FCD) across cohorts using a sliding-window approach.

#### 2.3.2. Fluidity of Functional Connectivity Dynamics

To characterize the temporal stability of the connectome, we computed FCD using a sliding-window correlation approach (window length = 20s, overlap = 1s). FCD fluidity was defined as the variance of the upper triangular elements (excluding the diagonal) of the time-resolved functional connectivity matrices (Lavanga et al., 2023; Hashemi et al., 2025). High fluidity reflects flexible transitions between segregated and integrated network states, whereas low fluidity indicates a more rigid functional architecture. To ensure comparability across subjects and independence from acquisition duration, we standardized the length of all time series prior to FCD computation. Scans shorter than 2.5 min were excluded from this specific analysis and scans exceeding this duration were truncated to this length.

### 2.4. Hierarchical Normative Modeling

Hierarchical normative modeling (Marquand et al., 2016, 2019) is a probabilistic framework that estimates individual deviations relative to a population distribution while explicitly modeling group-level variability (e.g., site or scanner effects). Such deviations can be interpreted as individualized biomarkers of disorders, and this framework provides a natural approach for modeling multiple levels of variation in heterogeneous data. In the following sections, we briefly present the general formulation of the model, clarify the incorporation of random effects and their structure, and describe the procedures for model selection and implementation.

#### 2.4.1. Mathematical Formulation

To model the changing distribution of functional connectivity across the lifespan, we employed a Generalized Additive Model for Location, Scale, and Shape (GAMLSS) (Stasinopoulos and Rigby, 2007; Dinga et al., 2021). GAMLSS provides a robust and flexible framework for modeling non-linear growth trajectories, as recommended by the World Health Organization (Borghi et al., 2006), and has been widely applied to estimate age-related trends from large datasets (Bethlehem et al., 2022; Sun et al., 2025). We utilized the Sinh-Arcsinh (SHASH) distribution (Jones and Pewsey, 2009) to accommodate non-Gaussian features of data. This flexible family is parameterized by location *µ*, which governs the central tendency; scale *σ*, which controls dispersion; skewness *ν*, which modulates asymmetry; and tail heaviness *τ*, which determines the weight of the distribution tails relative to a Gaussian. Note that when *ν* = 0 and *τ* = 1, the SHASH distribution is equivalent to a Gaussian. The connectivity metric value *y_ij_* for subject *i* in dataset or atlas *j* at age *x* is modeled directly by its conditional distribution:

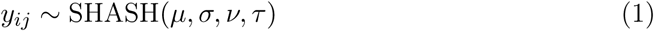

Rather than assuming a fixed functional form, we systematically constructed each parameter as a combination of fixed population-level trajectories and group-specific random effects. We define the generalized linear predictor for any distribution parameter *θ* ∈ {*µ, σ, ν, τ* } as:

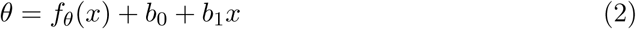

Here, *f_θ_*(*x*) represents the fixed-effect aging trajectory for parameter *θ*. Depending on the specific model, this trajectory can be a constant baseline, a linear age effect (denoted by the subscript *L*; *θ_L_* = *β*_0_ + *β*_1_*x*), or a penalized B-spline formulation (denoted by the subscript *S*.).

To accommodate technical heterogeneity, we introduce additive random effects. The term *b*_0_ represents a parameter-specific random intercept, while *b*_1_*x* represents a parameter-specific random slope on age. While our framework evaluated the inclusion of fixed age trends and random slopes across all higher-order shape parameters (*ν, τ*), components that suffered from severe predictive penalties or numerical instability are omitted from the final model notation. As an example, a model featuring linear maturation, dataset-specific baselines, and varying noise floors across sites is concisely denoted as SHASH(*µ_L_* +*b*_0_*, σ* +*b*_0_*, ν, τ*). A graphical illustration of such a model is given in Figure 3.

**Figure 3.**
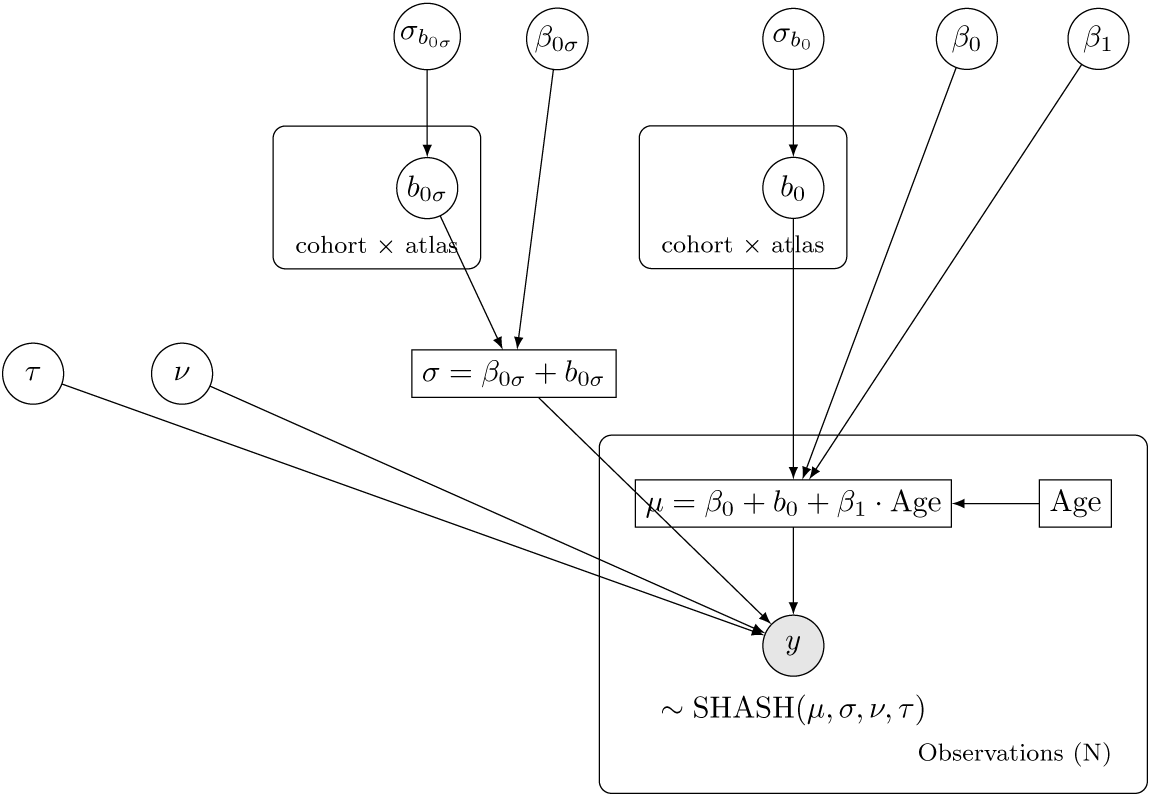
Graph representation of a Bayesian hierarchical mixed-effects model. A simple model featuring a SHASH likelihood, with a linear trend and cohort/atlas-specific baselines for the trend *µ* and the noise *σ* (random effects on the intercepts). In our notation, this model is denoted: SHASH(*µ_L_* + *b*_0_*, σ* + *b*_0_*, ν, τ*). Random variables to be sampled are represented as circles, while deterministic terms are squares. The frames are plates which mean that variables inside are repeated for every item in the group.

In the context of Bayesian hierarchical modeling, parameters must be assigned prior distributions, which will then be updated under Bayes’ rule with respect to the data. When parameters are defined as strictly fixed, default priors are set to *β*_0_ ∼ N (0, 5) for intercepts and *β*_1_ ∼ N (0, 10) for slopes. However, when random effects are imposed on a parameter (whether intercept or slope), the parameter is modeled as a mixed effect with its global fixed baseline tightened to N (0, 2). When non specified as being linear, higher-order fixed parameters were explicitly set to *ν* ∼ N (0, 1) and *τ* ∼ N (1, 1). To ensure the model remains identifiable, random effect offsets are drawn from a custom zero-sum Normal distribution. These group offsets are then scaled by a strictly positive standard deviation parameter modeled as *σ_b_* ∼ softplus(N (1, 1)).

#### 2.4.2. Interpretation of Random Effects

In the context of multi-site and multi-atlas normative modeling, each parameter-specific component of the random-effects structure targets a distinct source of observed methodological variability:

- Random intercepts on location (*µ*): This term (*b*_0_ on *µ*) captures systematic baseline offsets in connectivity magnitude. Including it allows the model to account for the fact that different parcellations (e.g., Schaefer vs. AAL) or preprocessing pipelines may yield consistently higher or lower average connectivity values, without imposing constraints on the shape of the developmental curve.
- Random intercepts on scale (*σ*): This term (*b*_0_ on *σ*) captures dataset-specific heteroscedasticity. It accommodates the likelihood that the precision of the data varies across domains. For instance, functional atlases may capture a wider range of inter-individual variability (higher variance) than structural ones, or specific scanners may introduce higher noise floors, independent of the mean signal strength.
- Higher-order parameters (*ν, τ*): Placing random effects on skewness (*ν*) or kurtosis (*τ*) parameters accounts for distributional artifacts. This accommodates scenarios where the shape of the population distribution itself (e.g., the presence of heavy tails or outliers) is driven by acquisition artifacts or processing strategies rather than biological reality.
- Random slopes (age interaction): This term (*b*_1_*x*) models the interaction between biology and methodology. By allowing the coefficient of age to vary by dataset or atlas, this term tests whether the observed rate of biological aging differs depending on the measurement tool.

#### 2.4.3. Random Effects Specification

While the general formulation defines the types of random effects applied to the parameters, we must also specify the grouping factors over which these effects vary. Given the sparsity of our aggregated data structure (where most datasets were represented in only one atlas space), the interaction term between “dataset” and “parcellation” (i.e., a dataset-specific atlas effect) was statistically unidentifiable for the majority of the cohort. We therefore specified the random-effects structure as additive across the two independent grouping factors. This specification assumes that the variance attributable to the parcellation scheme (driven by parcel homogeneity) and the variance attributable to the site (driven by scanner and protocol) are cumulative. While we validated the stability of this additive assumption within the fully crossed HCP sub-samples (Young Adult vs. Aging cohorts), we acknowledge that this model cannot capture potential idiosyncratic interactions between specific pipelines and atlas geometries in the non-crossed legacy datasets.

#### 2.4.4. Model Selection

In the main part of the study, we use a B-spline GAMLSS architecture with random intercepts on the location and scale parameters: SHASH(*µ_S_*+ *b*_0_*, σ* + *b*_0_). This formulation aligns with the current gold standard in large-scale normative modeling (Bethlehem et al., 2022; Sun et al., 2025) which assumes non-linear biological trajectories and addresses site-specific technical variance via static hierarchical offsets. However, to critically validate the assumptions underlying these multi-cohort charts, we performed a post-hoc model comparison evaluating the necessity of non-linear age terms and the potential existence of site-specific aging rates (random slopes).

Model comparison was performed using the Expected Log Pointwise Predictive Density (ELPD; Gelman et al. (2014)), estimated via Leave-One-Out Cross-Validation (LOO-CV) using Pareto-Smoothed Importance Sampling (PSIS-LOO; Vehtari et al. (2024)). For more details, see (Vehtari et al., 2017). Unlike traditional information criteria which rely on point estimates, ELPD-LOO provides a robust Bayesian estimate of out-of-sample predictive accuracy (Hashemi et al., 2021; Baldy et al., 2024). A model was considered superior if the improvement in ELPD was at least two times the standard error of the difference (|ΔELPD| *>* 2 × SE(Δ)).

We also report indicators of goodness-of-fit of the selected model, such as the coefficient of determination *R*^2^ of the selected model and the correlation *ρ* between the observed data and the predictions of the model.

#### 2.4.5. Software and Implementation

All normative modeling analyses were performed in Python 3.11 using the Predictive Clinical Neuroscience Toolkit (PCNtoolkit v1.1; Rutherford et al. (2023); Rutherford and Marquand (2025)). The toolkit was utilized to estimate the Bayesian GAMLSS models, providing a unified framework for fitting the SHASH likelihoods with penalized B-spline bases. Model diagnostics, including the computation of ELPD and Pareto-k values for PSIS-LOO cross-validation, were performed using ArviZ (Kumar et al., 2019).

## 3. Results

### 3.1. Trajectories of Human Brain Functional Connectome

First, we constructed lifespan trajectories of brain functional connectivity using standard metrics of static FC and dynamic FCD (see subsection 2.3). We modeled lifespan trajectories using penalized B-splines to characterize the baseline biological signal of functional aging without imposing constraints on the developmental curve. This flexible approach allowed us to detect potential non-linear phases, such as developmental peaks or accelerated decline in senescence, that a linear fit would otherwise mask.

To visualize the trajectories of brain functional connectivity within a unified reference frame, we harmonized the test data from all disparate cohorts into the common baseline space of the Human Connectome Project Young Adult (HCP-YA) dataset and Schaefer parcellation (Figure 4). Because our hierarchical modeling framework explicitly quantified the systematic shifts introduced by varying sites, scanners, and parcellation schemes as random effects, we used these learned estimates as statistical conversion factors. This post-hoc alignment removes the visual fragmentation caused by batch effects, allowing the underlying continuous biological pattern of functional aging to be visualized and interpreted as a coherent lifespan trajectory.

**Figure 4.**
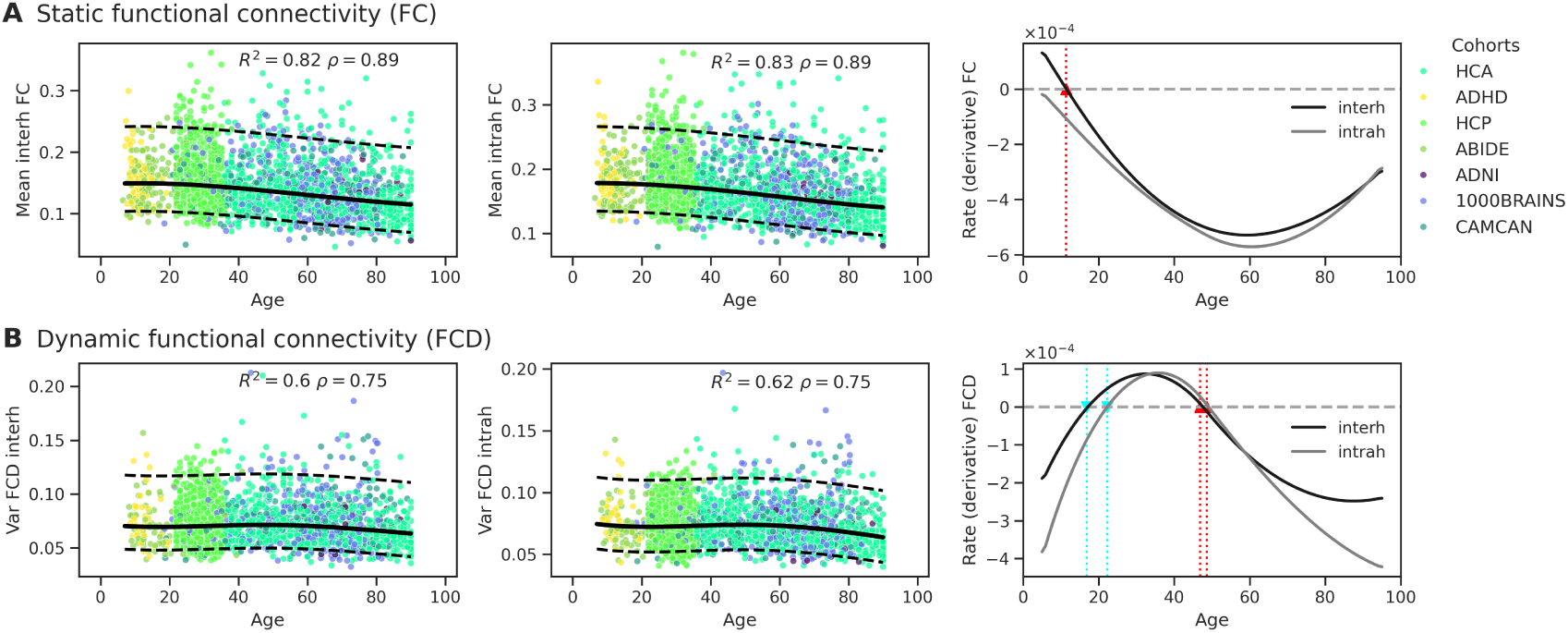
The lifespan normative trajectories of static (**A**) and dynamic (**B**) functional connectivity. Harmonized test data (dots, colored by cohort) is plotted along the normative trajectories fitted by the GAMLSS model (median, solid lines) and their 5% and 95% quantiles (dashed lines). The right panels show the rate of change (first derivative) of the normative curves for inter-(black) and intra-(grey) hemispheric connectivity. Zero-crossings of the derivative characterize the optima of the corresponding normative trajectory: minima are shown in cyan and maxima in red.

#### Static Functional Connectivity (FC)

The spline-based normative models shaped the evolution of global functional connectivity strength as a relatively stable decreasing trajectory across the adult lifespan following a peak in adolescence (see Figure 4). This trajectory of functional disintegration was pervasive, characterizing both long-range interhemispheric and local intra-hemispheric networks. While the rate of decline appeared conserved across these spatial scales, a distinct baseline offset was observed, with intrahemispheric edges exhibiting consistently higher connectivity magnitudes than interhemispheric coupling throughout the aging process. The absence of significant inflection points suggests that, for static connectivity, the aging process in this network is not characterized by abrupt phase transitions but rather by a cumulative biological decline. The shape of the first derivative would suggest a sustained cumulative decline throughout most of the lifespan, with slight deceleration of the decline rate after 60 years. Although this deceleration in old age is consistent with previously reported normative charts (Sun et al., 2025), their analysis revealed a lifespan trajectory peaking around age 40, contrary to our lifetime declining model.

#### Functional Connectivity Dynamics (FCD)

Beyond static coupling, we quantified the temporal stability of the connectome using the FCD measure, capturing the variance of time-resolved correlation matrices across sliding windows as a proxy for macroscopic network fluidity. By expanding our normative framework to include FCD, our spline models (see Figure 4) revealed a complex, non-monotonic developmental trajectory. From childhood to early adulthood (approximately ages 18–20), macroscopic fluidity exhibits a continuous but decelerating decrease, reaching a local minimum (indicated by the zero-crossing of the first derivative). This phase appears to be delayed by 2–4 years for intra-hemispheric connections compared to inter-hemispheric fluidity; however, it remains to be explored whether this delay is statistically significant. Crossing into early adulthood, the derivative of the FCD curve becomes positive and accelerates until reaching an inflection point in the third decade of life, driving a prolonged expansion of network fluidity that peaks around age 50. From age 50 onward, the derivative crosses zero again and becomes progressively more negative, marking a terminal decline in fluidity.

#### Cohort-Specific Trajectories and Cross-Atlas Projections

The predictions of the model at the level of individual cohorts and parcellation schemes are detailed for the mean interhemispheric FC, and variance FCD in Figure 5. The solid regression lines and dotted normative quantiles represent the specific GAMLSS estimates derived for each dataset-atlas combination. As evidenced by the distribution of the test data (grey points), the hierarchical architecture with random intercepts successfully calibrates to the distinct baseline connectivity levels and noise profiles of each cohort. Crucially, because the framework explicitly parameterizes the brain atlas as a distinct random effect, it enables the translation of data across methodological space. To demonstrate this, we projected subjects into parcellation schemes in which they were never actually processed (Figure 5, green points). By extracting a cohort’s baseline biological variance and applying the model’s learned atlas-specific offsets, the model projects subjects into unobserved spatial configurations. Similar plots for the intra-hemispheric measures are shown in Supplementary Figure S1.

**Figure 5.**
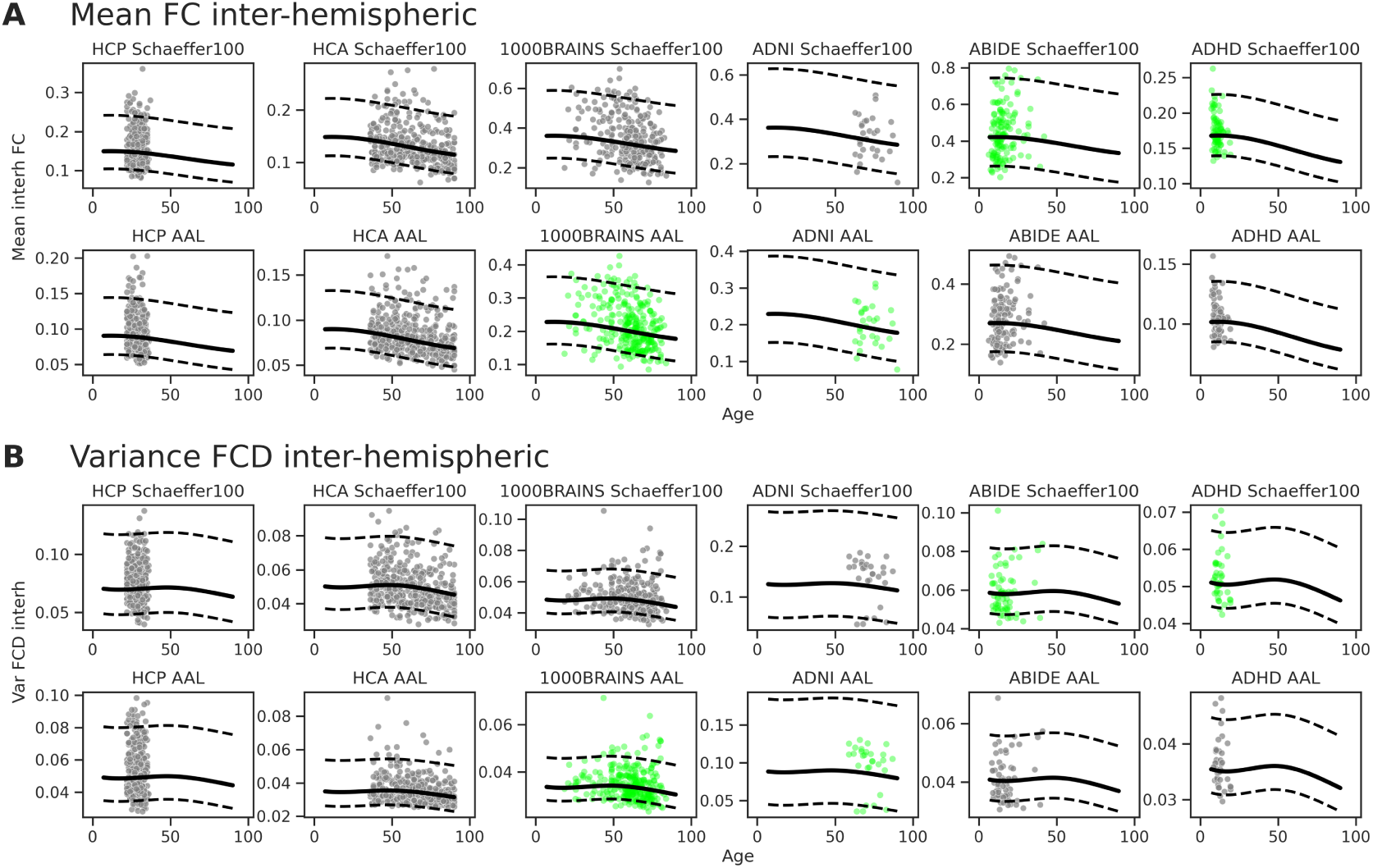
Cohort-specific normative trajectories and cross-atlas projections for (**A**) the mean of interhemispheric FC and (**B**) the variance of inter-hemispheric FCD. Solid lines and dotted boundaries represent the cohort- and atlas-specific median regression lines and normative quantiles estimated by the hierarchical GAMLSS model. Test data is shown in grey; the cross-atlas projections (in green) of test data for missing cohort-atlas combinations are generated by leveraging the model’s estimated random effects.

### 3.2. Sensitivity to Connectivity Metrics

We evaluated the robustness of normative trajectories across five additional metrics of functional coupling (Figure 6). Among correlation-based measures, Pearson correlation, Spearman’s *ρ*, and mutual information all exhibited robust, reliable trajectories with high goodness-of-fit (*R*^2^ ≈ 0.8). These metrics showed a consistent, monotonic pattern of age-related decline, indicating that the signal of functional disintegration is well-preserved across linear, rank-based, and information-theoretic measures. Magnitude-squared coherence, which isolates coupling within the low-frequency fluctuation range (0.01–0.1 Hz), mirrored the behavior of the time-domain metrics. It exhibited a robust age-related decline (despite a lower *R*^2^ = 0.75), confirming that the loss of functional integration is observable in the frequency domain as well as in the time domain. The precision metric (partial correlation) performed the worst, paradoxically. The models frequently encountered numerical convergence warnings, and the estimated trajectories were essentially flat (*R*^2^ ≈ 0.92 but with a negligible slope), suggesting that precision matrices are too sensitive to noise or sample size variations to serve as a robust normative biomarker in heterogeneous aggregate datasets.

**Figure 6.**
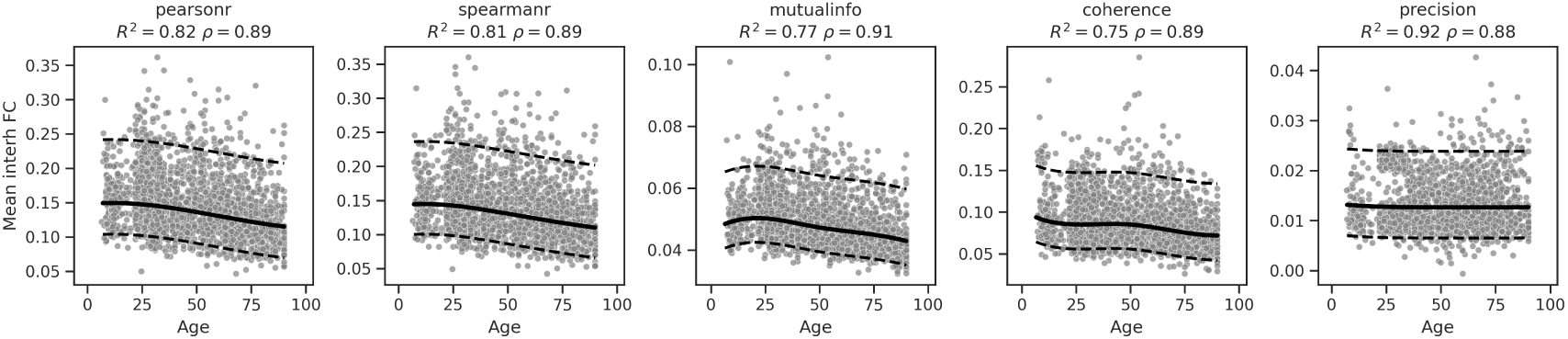
Normative trajectories of static functional connectivity computed according to several metrics from standard statistics (Pearson and Spearman’s correlation coefficients, precision matrix), information theory (mutual information) and spectral analysis (magnitude-squared coherence). Normative trajectory (median) and quantiles (5% and 95%) are shown in black solid and dashed lines, while test data is shown in grey.

### 3.3. Quantification of Extreme Deviations

To evaluate the out-of-sample calibration and clinical sensitivity of the normative model, we quantified the proportion of extreme deviations (|*Z*| *>* 1.645, corresponding to the 10% theoretical expectation) across the healthy and diagnosed test sets.

First, we assessed the absolute calibration of the normative model on the healthy control data. Overall, the globally aggregated control cohort robustly approximated the expected 10% deviation rate. To rigorously test whether local site-specific variance profiles skewed this calibration, we performed exact binomial tests comparing the proportion of extreme deviations in each local control group against the theoretical 10% expectation. After Benjamini-Hochberg False Discovery Rate (FDR) correction (Benjamini and Hochberg, 1995), local control baselines did not significantly differ from the 10% expectation in the vast majority of datasets and metrics (see Supplementary Table S2). The only statistically significant deviation was observed in the 1000BRAINS cohort for inter-hemispheric FCD measure. The general lack of significant deviation across the remaining disparate sites (including highly specific sub-cohorts such as ADHD-200 and ADNI) demonstrates that the hierarchical GAMLSS framework successfully accommodates multi-site heterogeneity. By modeling site and atlas as random effects, the framework aligns both the global mean trajectories and the local variance profiles without requiring aggressive and variance-destroying pre-harmonization.

Second, we evaluated the clinical sensitivity of these calibrated thresholds. We compared the proportion of extreme deviations between diagnosed cohorts and their local control counterparts using Fisher’s Exact Tests, adjusted for multiple comparisons via FDR. While qualitative numerical trends were present, such as an inflation of extreme deviations in Mild Cognitive Impairment (MCI) or ADHD relative to their respective local controls (see Supplementary Table S2 and Figure S3) no diagnostic group exhibited a statistically significant increase in extreme deviations after multiple comparison correction (all *p_F DR_ >* 0.05, Supplementary Table S2).

### 3.4. Model Comparison

Having characterized non-linear biological shapes of functional aging, we rigorously evaluated whether the additional complexity of the spline models was statistically justified for constructing a generalizable reference chart. We systematically evaluated candidate GAMLSS specifications to identify a generative model that balances biological flexibility with predictive robustness. Figure 7 shows Bayesian model comparison featuring the Expected Log Pointwise Predictive Density (ELPD) of statistical models, ranked based on their performance, for the inter-hemispheric static functional connectivity and fluidity. A similar figure for intra-hemispheric connectivities is given in the Supplementary Figure S2. Moreover, Supplementary Table S1 reports all the top-performing models across features and metrics in terms of their out-of-sample predictive accuracy. The choice of likelihood function proved to be the primary determinant of model performance. Models utilizing the flexible Sinh-Arcsinh (SHASH) distribution consistently outperformed Gaussian equivalents by a substantial margin of at least two standard deviations (see Figure 7, and Supplementary Table S1), highlighting the necessity of accounting for the non-Gaussian nature of functional connectivity metrics. Specifically, estimating higher-order moments (skewness *ν* and kurtosis *τ*) significantly improved model fit. However, introducing random effects on these shape parameters critically degraded predictive performance, suggesting that while the distribution shape varies between metrics, it remains relatively stable across sites and ages within a metric.

**Figure 7.**
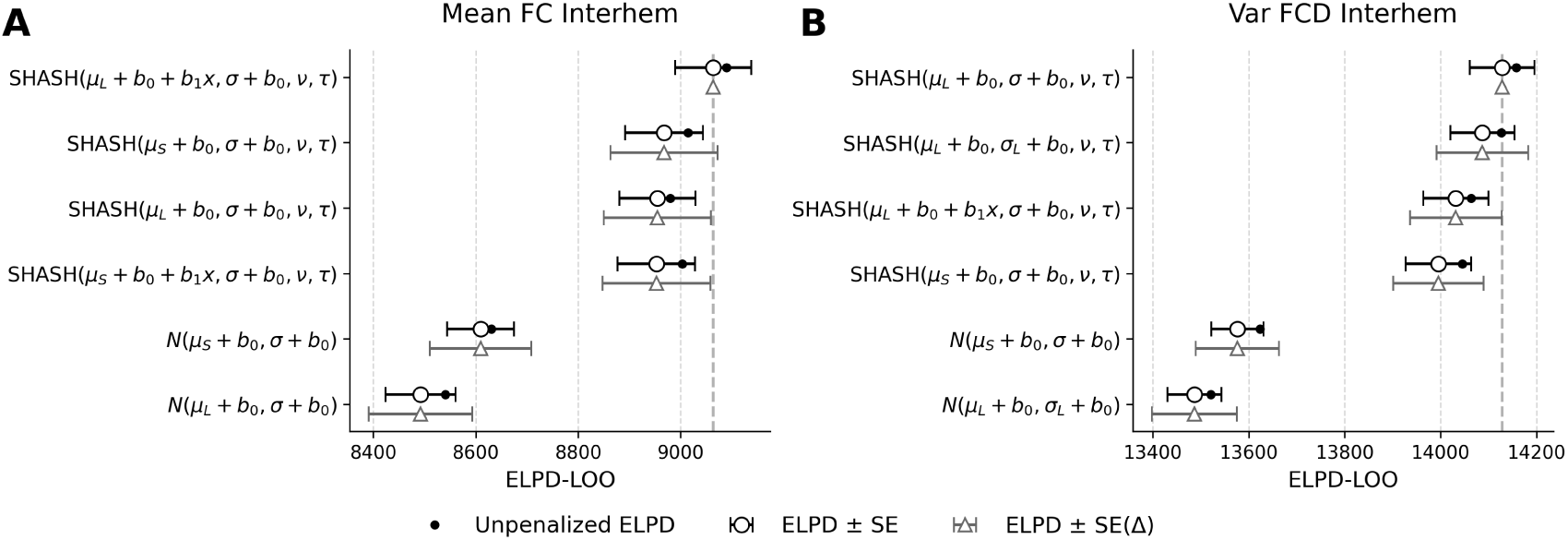
Bayesian model comparison for inter-hemispheric functional connectivity (**A**) and fluidity measured by variance of dynamic functional connectivity (**B**). Models are ranked on the y-axis according to their predictive accuracy, with the best-performing (reference) model placed at the top. The x-axis represents the Leave-One-Out Expected Log Pointwise Predictive Density (ELPD-LOO) as the measure of predictive accuracy using cross-validation. The vertical dashed grey line marks the ELPD of the reference model. Black circles indicate the absolute ELPD estimate for each model, with horizontal black error bars representing *±*1 standard error (SE) of that estimate. Grey triangles are plotted such that their visual distance from the vertical dashed line corresponds exactly to the difference in ELPD (Δ ELPD) compared to the top model. The horizontal grey error bars denote *±*1 standard error of this difference, *SE*(Δ). We consider models ranking within 2 *× SE*(Δ) of the top model to be statistically indistinguishable. Legend: *µ_L_* = Linear trend; *µ_S_* = Spline trend; *σ_L_* = Linear trend on scale; *b*_0_ = Parameter-specific random intercept; *b*_1_*x* = Parameter-specific random slope.

Our results indicate overwhelming support for including random intercepts on both the location *µ* and scale *σ* parameters (see Supplementary Table S1, where for any feature and metric, all models within two standard deviations of the top performer model were SHASH). This indicates that while different sites and parcellations introduce static offsets in mean connectivity and signal variance (heteroscedasticity), these effects can be effectively modeled as additive constants. Crucially, introducing random slopes for the age term did not significantly improve out-of-sample predictive accuracy. In many cases, the simpler random-intercept-only model was statistically indistinguishable from the complex model, suggesting that the trajectory of functional decline is largely conserved across heterogeneous acquisition protocols, even if the baseline values differ. Additionally, spline-based models did not yield a statistically significant improvement in predictive accuracy with respect to their linear counterparts.

## 4. Discussion

### 4.1. The Lifespan Trajectory of Fluidity

While static global connectivity exhibits a largely monotonic age-related decline, our extension of the normative framework to Functional Connectivity Dynamics (FCD) revealed a distinctly non-linear, triphasic trajectory. This highlights a profound biological decoupling between the structural integrity of the connectome and its temporal flexibility. The initial decrease in dynamic variance from childhood to early adulthood likely reflects the well-documented transition from a hyper-flexible, unconstrained pediatric brain into a mature, segregated, and highly efficient resting-state topology (Hutchison and Morton, 2015; Fair et al., 2009). As core networks, such as the default mode and frontoparietal networks, stabilize during adolescence, the system’s random temporal fluctuations are minimized (López-Vicente et al., 2021). Strikingly, crossing into mid-adulthood, network fluidity expands, peaking around age 50. We interpret this mid-life surge as the acquisition of peak metastability—an expanded dynamic repertoire that allows the brain to rapidly transition between distinct cognitive states (Deco et al., 2017; Garrett et al., 2013). This suggests that while absolute structural and static connection strength may begin degrading in early adulthood, the dynamic flexibility of the system is actively preserved and matures to support the complex cognitive switching demands of middle age. Finally, the terminal decline in fluidity beyond age 50 marks the onset of senescent rigidity. In this phase, the age-related breakdown of structural integrity and neurotransmitter signaling outpaces compensatory mechanisms, leading to a loss of network fluidity. The connectome becomes increasingly rigid, trapping the brain in fewer, less flexible network states, which closely aligns with the cognitive inflexibility characteristically observed in advanced aging (Battaglia et al., 2020).

### 4.2. Biological Invariance and the Stability of Aging Trajectories

Beyond the selection of the normative reference model, the comparative performance of the candidate specifications offers two significant biological insights into the nature of functional connectivity aging. First, the observation that random slopes for age did not improve predictive accuracy has important implications for multi-site neuroimaging. According to the “maximal random-effects structure” recommended by Barr et al. (2013), the omission of random slopes is justified only when the variance of those slopes is negligible. The fact that model comparison favored static slopes suggests that the normative aging trajectory of functional connectivity in this metric is largely monotonic and conserved across acquisition sites, with little support for site-specific rates of decline. While scanners and parcellations introduce significant static offsets (evidenced by the strong support for random intercepts), they do not appear to systematically alter the estimated aging trajectory.

Second, the near-equivalence of the linear and spline-based models indicates that, within the studied age range, the decline in functional connectivity follows a largely monotonic and stable course. While we retained the spline model to robustly accommodate potential edge-case non-linearities, the lack of a strong predictive penalty for the linear constraint suggests that the aging process in this network is not characterized by abrupt phase transitions as in complex systems, but rather by a steady, cumulative biological weathering. However, beyond a purely biological interpretation, the statistical preference for a linear trajectory must be contextualized within the fragmented structure of our aggregated data. Our cohort is a mosaic of diverse datasets, several of which occupy restricted, non-overlapping windows of the human lifespan (e.g., ADHD-200 in childhood, ADNI in late senescence). In such a piecewise landscape, highly flexible non-linear models like B-splines are susceptible to overfitting; they may misinterpret local, dataset-specific scanner noise or sampling biases at specific age brackets as true biological inflection points. Therefore, the near-equivalence of the linear model does not necessarily preclude the existence of highly subtle, non-linear biological phases; rather, it indicates that a linear constraint acts as a powerful regularizer (Nozari et al., 2024). It yields the most robust, generalizable summary of functional decline across disparate sites. Crucially, the selection of this linear baseline is not merely an artifact of the model failing to accommodate divergent site trajectories. According to the “maximal random-effects structure”, if the underlying rates of aging were genuinely heterogeneous across acquisition protocols, the model selection would have strongly favored the inclusion of random slopes for age.

### 4.3. Methodological Considerations and Hierarchical Structure

A notable methodological consideration in our framework is the resolution of the random effects structure. Previous normative modeling efforts, including the structural (Bethlehem et al., 2022) and the functional (Sun et al., 2025) brain charts, explicitly modeled batch effects at the micro-level of individual acquisition sites and scanners. In contrast, our study defined random effects at the macro-level of the aggregated datasets without nesting individual intra-cohort sites.

Our approach represents a deliberate trade-off between granular variance partitioning and model stability. The GAMLSS architecture is computationally demanding when using gradient-based Hamiltonian Monte Carlo for unbiased sampling. Introducing deeply nested hierarchies across highly heterogeneous datasets frequently leads to over-parameterization and convergence failures, especially when some acquisition sites contain very few subjects or span narrow age ranges. By restricting our random intercepts to the cohort level, we maintained a stable, universal generative model capable of converging across the entire metric landscape. Importantly, this macro-level approach does not simply ignore intra-cohort variance. Because our framework explicitly models the scale parameter, unmodeled inter-site discrepancies within aggregated consortia are naturally captured as increased heteroscedasticity in those cohorts.

### 4.4. Clinical Sensitivity and the Limits of Global Scalars

A critical evaluation of any normative chart requires assessing its calibration and sensitivity to pathological deviations. Applying our harmonized consensus model to compute individualized Z-scores revealed a good baseline calibration: in the vast majority of local healthy control cohorts, the empirical rate of extreme deviations aligned closely with the theoretical 10% expectation (Supplementary Table S2). This confirms that the hierarchical harmonization successfully absorbed site- and cohort-specific technical variance, preventing the artificial inflation of outlier rates that plagues poorly harmonized multi-site data. However, despite this accurate baseline calibration, we observed no statistically significant inflation of extreme deviations in diagnostic groups (e.g., ADHD, AD, MCI) relative to their local controls. While initially counterintuitive, this highlights a fundamental biological limitation of macroscopic, averaged whole-brain metrics. Such global summaries collapse the rich spatial heterogeneity of brain organization into a single scalar, potentially masking localized alterations that characterize many neurological and psychiatric conditions. Detecting subtle pathological deviations therefore require more heterogeneous and spatially resolved metrics that preserve regional variability, network-specific interactions, or edge-level connectivity patterns rather than relying on whole-brain averages (Zalesky et al., 2010; Faskowitz et al., 2020; Wolfers et al., 2018; Segal et al., 2023; Dahmen et al., 2025; Hashemi et al., 2025).

## Code availability

All code is available on GitHub (https://github.com/ins-amu/NormativeModeling) and in Ebrains collaboratory drive task 3.3.

## Data availability

Data were provided in part by the Human Connectome Project, WU-Minn Consortium (Principal Investigators: David Van Essen and Kamil Ugurbil; 1U54MH091657) funded by the 16 NIH Institutes and Centers that support the NIH Blueprint for Neuroscience Research; and by the McDonnell Center for Systems Neuroscience at Washington University. Data in this publication were provided in part by the Aging Adult Vulnerability and Resiliency in the Aging Adult Brain Connectome (AABC) project (U19AG073585) and the Human Connectome Project in Aging (HCP-A, U01AG052564) funded by the National Institute on Aging (NIA).

Data collection and sharing for this project was provided by the Cambridge Centre for Ageing and Neuroscience (CamCAN). CamCAN funding was provided by the UK Biotechnology and Biological Sciences Research Council (grant number BB/H008217/1), together with support from the UK Medical Research Council and University of Cambridge, UK. Data used in this work were obtained from the CamCAN repository (https://cam-can.mrc-cbu.cam.ac.uk/dataset/).

The ADHD-200 data used in this study were provided by the ADHD-200 Consortium and the International Neuroimaging Data-sharing Initiative (INDI). Preprocessed data were obtained from the ADHD-200 Preprocessed repository, provided freely by The Neuro Bureau and the Preprocessed Connectomes Project (PCP). We acknowledge the contributing sites and the developers of the preprocessing pipelines and imaging analysis packages utilized by The Neuro Bureau. The ABIDE data used in this study were provided by the Autism Brain Imaging Data Exchange (ABIDE) Consortium and the International Neuroimaging Data-sharing Initiative (INDI). Preprocessed functional and structural data were provided by the Preprocessed Connectomes Project (PCP). Primary support for the work by ABIDE was provided by the NIMH (K23MH087770 and R03MH096321) and the Leon Levy Foundation.

Data used in the preparation of this article were obtained from the Alzheimer’s Disease Neuroimaging Initiative (ADNI) database (adni.loni.usc.edu). The ADNI was launched in 2003 as a public-private partnership, led by Principal Investigator Michael W. Weiner, MD. The primary goal of ADNI has been to test whether serial magnetic resonance imaging (MRI), positron emission tomography (PET), other biological markers, and clinical and neuropsychological assessment can be combined to measure the progression of mild cognitive impairment (MCI) and early Alzheimer’s disease (AD). Data collection and sharing for the Alzheimer’s Disease Neuroimaging Initiative (ADNI) is funded by the National Institute on Aging (National Institutes of Health Grant U19AG024904). The grantee organization is the Northern California Institute for Research and Education. In the past, ADNI has also received funding from the National Institute of Biomedical Imaging and Bioengineering, the Canadian Institutes of Health Research, and private sector contributions through the Foundation for the National Institutes of Health (FNIH) including generous contributions from the following: AbbVie, Alzheimer’s Association; Alzheimer’s Drug Discovery Foundation; Araclon Biotech; BioClinica, Inc.; Biogen; Bristol-Myers Squibb Company; CereSpir, Inc.; Cogstate; Eisai Inc.; Elan Pharmaceuticals, Inc.; Eli Lilly and Company; EuroImmun; F. Hoffmann-La Roche Ltd and its affiliated company Genentech, Inc.; Fujirebio; GE Healthcare; IXICO Ltd.; Janssen Alzheimer Immunotherapy Research & Development, LLC.; Johnson & Johnson Pharmaceutical Research &Development LLC.; Lumosity; Lundbeck; Merck & Co., Inc.; Meso Scale Diagnostics, LLC.; NeuroRx Research; Neurotrack Technologies; Novartis Pharmaceuticals Corporation; Pfizer Inc.; Piramal Imaging; Servier; Takeda Pharmaceutical Company; and Transition Therapeutics.

## Acknowledgements

This research has received funding from EU’s Horizon 2020 Framework Programme for Research and Innovation under the Specific Grant Agreements No. 101147319 (EBRAINS 2.0 Project), No. 101137289 (Virtual Brain Twin Project), and government grant managed by the Agence Nationale de la Recherche reference ANR-22-PESN-0012 (France 2030 program). The funders had no role in study design, data collection and analysis, decision to publish, or preparation of the manuscript.

## Author contributions

Conceptualization: S.P., M.H., and V.J. Methodology: N.B., P.T, S.P., M.H., and V.J. Software: N.B, and M.H. Data curation: N.B, and P.T. Investigation: N.B. Visualization: N.B. Supervision: M.H., and V.J. Funding acquisition: M.H., and V.J. Writing - original draft: N.B., and M.H., and V.J. Writing - review & editing: N.B, P.T, S.P., M.H., and V.J.

## 5. Supplementary

**Figure S1.**
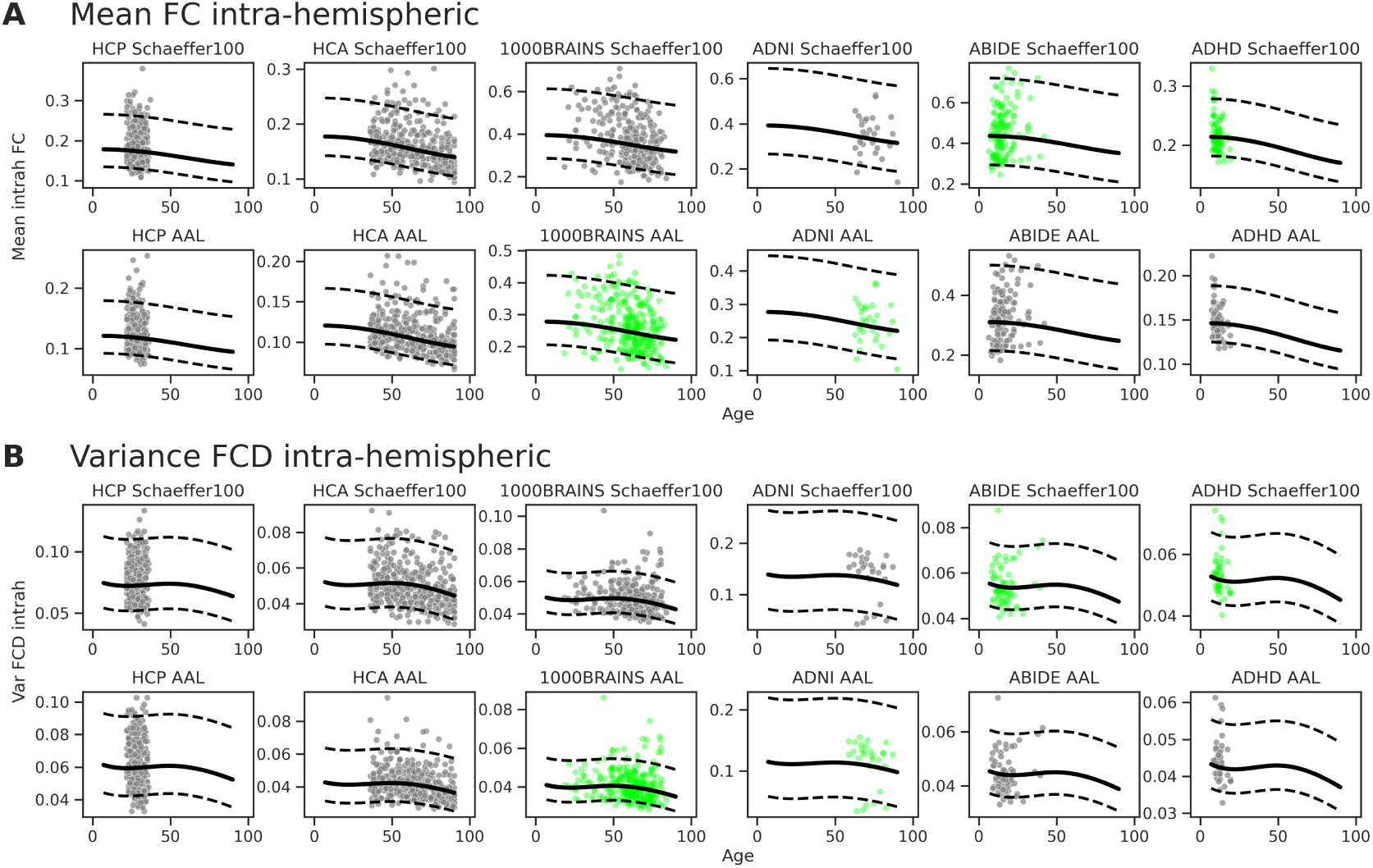
Cohort-specific normative trajectories and cross-atlas projections for (**A**) FC intrahemispheric and (**B**) FCD intra-hemispheric. Solid lines and dotted boundaries represent the cohort- and atlas-specific median regression lines and normative quantiles estimated by the hierarchical GAMLSS model. Test data is shown in grey; the cross-atlas projections (in green) of test data for missing cohort-atlas combinations are generated by leveraging the model’s estimated random effects.

**Figure S2.**
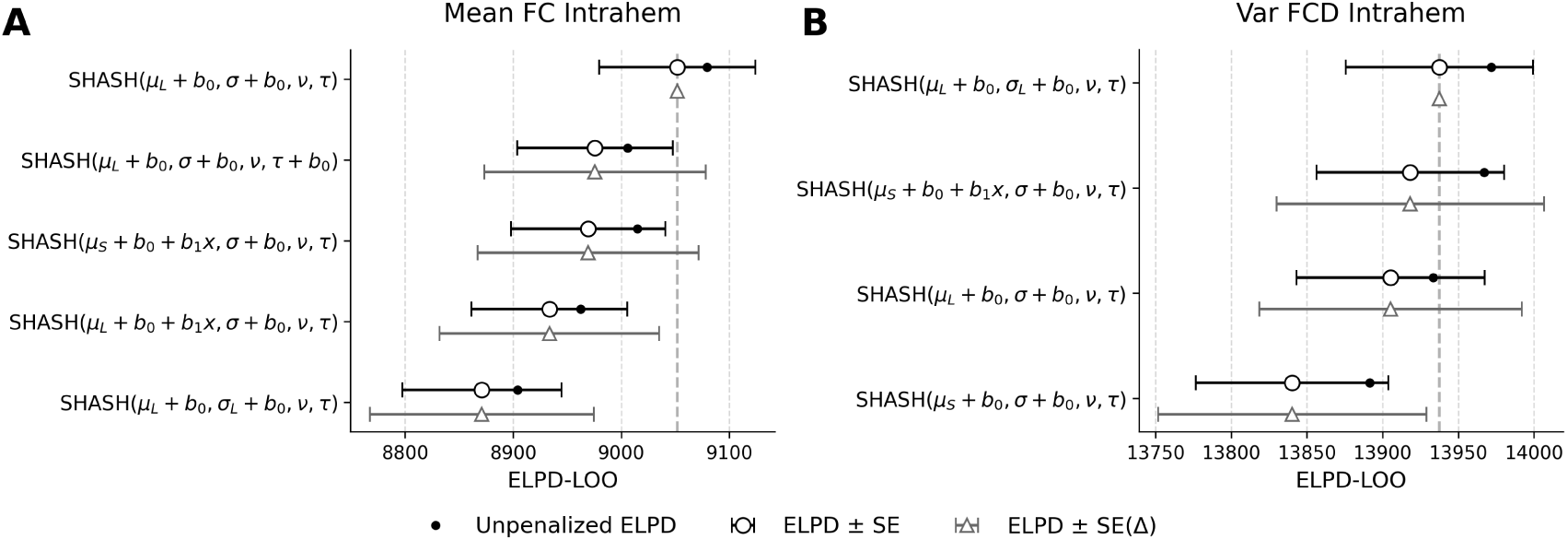
Bayesian model comparison for intrahemispheric functional connectivity (**A**) and fluidity via variance of dynamic functional connectivity (**B**). Models are ranked on the y-axis according to their predictive accuracy, with the best-performing (reference) model placed at the top. The x-axis represents the Leave-One-Out Expected Log Pointwise Predictive Density (ELPD-LOO), as the measure of predictive accuracy using cross-validation. The vertical dashed grey line marks the ELPD of the reference model. Black circles indicate the absolute ELPD estimate for each model, with horizontal black error bars representing *±*1 standard error (SE) of that estimate. Grey triangles are plotted such that their visual distance from the vertical dashed line corresponds exactly to the difference in ELPD (Δ ELPD) compared to the top model. The horizontal grey error bars denote *±*1 standard error of this difference, *SE*(Δ). We considers models ranking within 2 *× SE*(Δ) of the top model as statistically indistinguishable from it. Legend: *µ_L_* = Linear trend; *µ_S_* = Spline trend; *σ_L_* = Linear trend on scale; *b*_0_ = Parameter-specific random intercept; *b*_1_*x* = Parameter-specific random slope.

**Figure S3.**
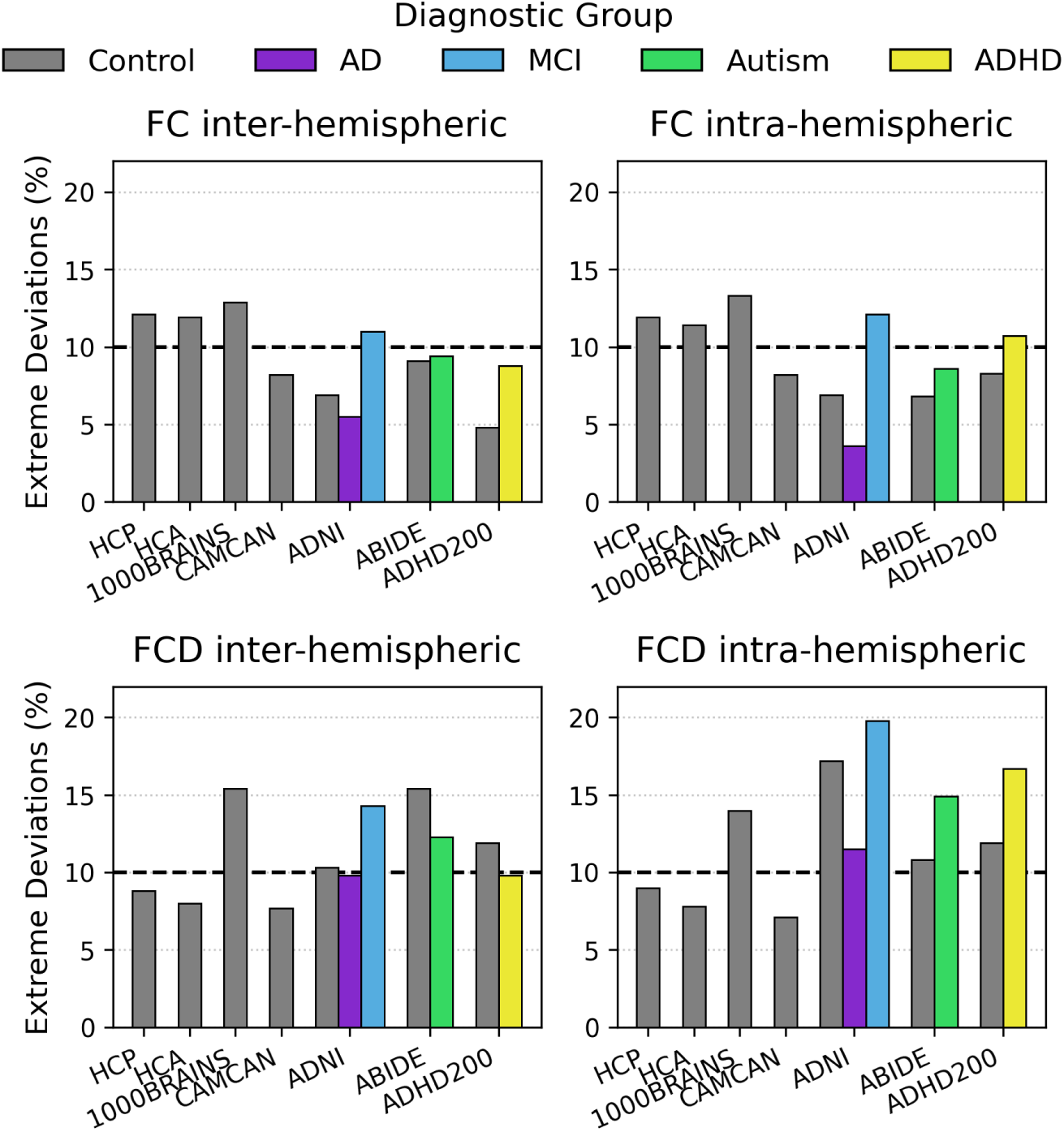
Percentage of extreme deviations (*|Z| >* 1.645) across healthy controls and clinical cases for static and dynamic functional connectivity (denoted by FC and FCD, respectively). The theoretical baseline is 10% (dashed black line).

**Table S1.**
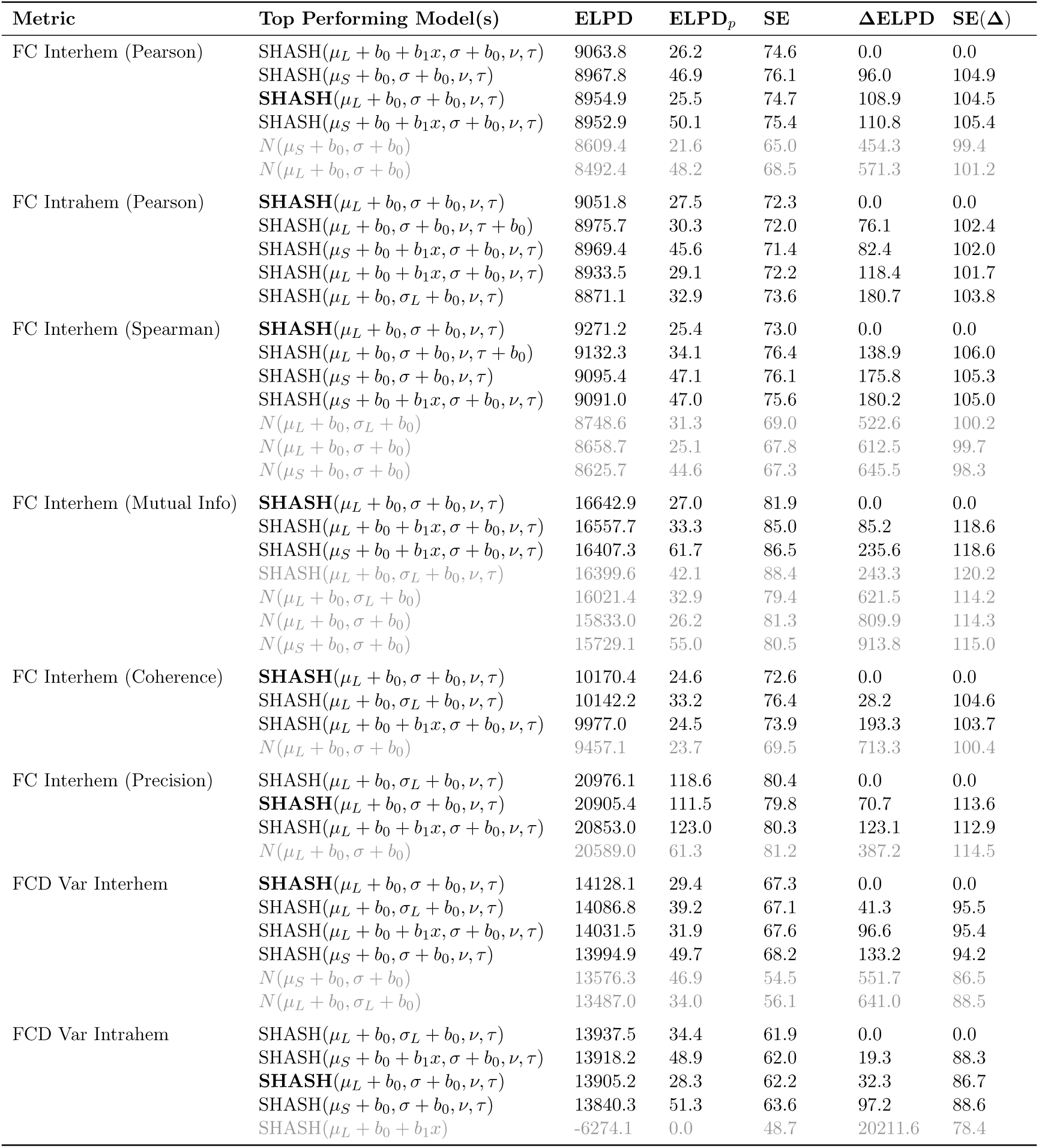
Best statistical model specifications ranked by predictive accuracy: Expected Log Pointwise Predictive Density **(ELPD)**, Estimated effective number of parameters (**ELPD***_p_*), Standard Error of the ELPD (**SE**), Difference of ELPD from best model (**Δ**ELPD) and Standard Error of the difference (SE(**Δ**)).Models within 2 standard errors SE(**Δ**) of the top-performing model are in black, other in gray. Only models that did not encounter warnings were reported^*^. The selected common reference model is shown in bold. Legend: *µ_L_* = Linear trend; *µ_S_* = Spline trend; *σ_L_* = Linear trend on scale; *b*_0_ = Parameter-specific random intercept; *b*_1_*x* = Parameter-specific random slope. ^*^ Except for FC Interhem (Precision) where all the models had warnings.

| Metric | Top Performing Model(s) | ELPD | ELPD<sub>p</sub> | SE | ΔELPD | SE(Δ) |
| --- | --- | --- | --- | --- | --- | --- |
| FC Interhem (Pearson) | SHASH($\mu_L + b_0 + b_1x, \sigma + b_0, \nu, \tau$) | 9063.8 | 26.2 | 74.6 | 0.0 | 0.0 |
| | SHASH($\mu_S + b_0, \sigma + b_0, \nu, \tau$) | 8967.8 | 46.9 | 76.1 | 96.0 | 104.9 |
| | **SHASH**($\mu_L + b_0, \sigma + b_0, \nu, \tau$) | 8954.9 | 25.5 | 74.7 | 108.9 | 104.5 |
| | SHASH($\mu_S + b_0 + b_1x, \sigma + b_0, \nu, \tau$) | 8952.9 | 50.1 | 75.4 | 110.8 | 105.4 |
| | $N(\mu_S + b_0, \sigma + b_0)$ | 8609.4 | 21.6 | 65.0 | 454.3 | 99.4 |
| | $N(\mu_L + b_0, \sigma + b_0)$ | 8492.4 | 48.2 | 68.5 | 571.3 | 101.2 |
| FC Intrahem (Pearson) | **SHASH**($\mu_L + b_0, \sigma + b_0, \nu, \tau$) | 9051.8 | 27.5 | 72.3 | 0.0 | 0.0 |
| | SHASH($\mu_L + b_0, \sigma + b_0, \nu, \tau + b_0$) | 8975.7 | 30.3 | 72.0 | 76.1 | 102.4 |
| | SHASH($\mu_S + b_0 + b_1x, \sigma + b_0, \nu, \tau$) | 8969.4 | 45.6 | 71.4 | 82.4 | 102.0 |
| | SHASH($\mu_L + b_0 + b_1x, \sigma + b_0, \nu, \tau$) | 8933.5 | 29.1 | 72.2 | 118.4 | 101.7 |
| | SHASH($\mu_L + b_0, \sigma_L + b_0, \nu, \tau$) | 8871.1 | 32.9 | 73.6 | 180.7 | 103.8 |
| FC Interhem (Spearman) | **SHASH**($\mu_L + b_0, \sigma + b_0, \nu, \tau$) | 9271.2 | 25.4 | 73.0 | 0.0 | 0.0 |
| | SHASH($\mu_L + b_0, \sigma + b_0, \nu, \tau + b_0$) | 9132.3 | 34.1 | 76.4 | 138.9 | 106.0 |
| | SHASH($\mu_S + b_0, \sigma + b_0, \nu, \tau$) | 9095.4 | 47.1 | 76.1 | 175.8 | 105.3 |
| | SHASH($\mu_S + b_0 + b_1x, \sigma + b_0, \nu, \tau$) | 9091.0 | 47.0 | 75.6 | 180.2 | 105.0 |
| | $N(\mu_L + b_0, \sigma_L + b_0)$ | 8748.6 | 31.3 | 69.0 | 522.6 | 100.2 |
| | $N(\mu_L + b_0, \sigma + b_0)$ | 8658.7 | 25.1 | 67.8 | 612.5 | 99.7 |
| | $N(\mu_S + b_0, \sigma + b_0)$ | 8625.7 | 44.6 | 67.3 | 645.5 | 98.3 |
| FC Interhem (Mutual Info) | **SHASH**($\mu_L + b_0, \sigma + b_0, \nu, \tau$) | 16642.9 | 27.0 | 81.9 | 0.0 | 0.0 |
| | SHASH($\mu_L + b_0 + b_1x, \sigma + b_0, \nu, \tau$) | 16557.7 | 33.3 | 85.0 | 85.2 | 118.6 |
| | SHASH($\mu_S + b_0 + b_1x, \sigma + b_0, \nu, \tau$) | 16407.3 | 61.7 | 86.5 | 235.6 | 118.6 |
| | SHASH($\mu_L + b_0, \sigma_L + b_0, \nu, \tau$) | 16399.6 | 42.1 | 88.4 | 243.3 | 120.2 |
| | $N(\mu_L + b_0, \sigma_L + b_0)$ | 16021.4 | 32.9 | 79.4 | 621.5 | 114.2 |
| | $N(\mu_L + b_0, \sigma + b_0)$ | 15833.0 | 26.2 | 81.3 | 809.9 | 114.3 |
| | $N(\mu_S + b_0, \sigma + b_0)$ | 15729.1 | 55.0 | 80.5 | 913.8 | 115.0 |
| FC Interhem (Coherence) | **SHASH**($\mu_L + b_0, \sigma + b_0, \nu, \tau$) | 10170.4 | 24.6 | 72.6 | 0.0 | 0.0 |
| | SHASH($\mu_L + b_0, \sigma_L + b_0, \nu, \tau$) | 10142.2 | 33.2 | 76.4 | 28.2 | 104.6 |
| | SHASH($\mu_L + b_0 + b_1x, \sigma + b_0, \nu, \tau$) | 9977.0 | 24.5 | 73.9 | 193.3 | 103.7 |
| | $N(\mu_L + b_0, \sigma + b_0)$ | 9457.1 | 23.7 | 69.5 | 713.3 | 100.4 |
| FC Interhem (Precision) | SHASH($\mu_L + b_0, \sigma_L + b_0, \nu, \tau$) | 20976.1 | 118.6 | 80.4 | 0.0 | 0.0 |
| | **SHASH**($\mu_L + b_0, \sigma + b_0, \nu, \tau$) | 20905.4 | 111.5 | 79.8 | 70.7 | 113.6 |
| | SHASH($\mu_L + b_0 + b_1x, \sigma + b_0, \nu, \tau$) | 20853.0 | 123.0 | 80.3 | 123.1 | 112.9 |
| | $N(\mu_L + b_0, \sigma + b_0)$ | 20589.0 | 61.3 | 81.2 | 387.2 | 114.5 |
| FCD Var Interhem | **SHASH**($\mu_L + b_0, \sigma + b_0, \nu, \tau$) | 14128.1 | 29.4 | 67.3 | 0.0 | 0.0 |
| | SHASH($\mu_L + b_0, \sigma_L + b_0, \nu, \tau$) | 14086.8 | 39.2 | 67.1 | 41.3 | 95.5 |
| | SHASH($\mu_L + b_0 + b_1x, \sigma + b_0, \nu, \tau$) | 14031.5 | 31.9 | 67.6 | 96.6 | 95.4 |
| | SHASH($\mu_S + b_0, \sigma + b_0, \nu, \tau$) | 13994.9 | 49.7 | 68.2 | 133.2 | 94.2 |
| | $N(\mu_S + b_0, \sigma + b_0)$ | 13576.3 | 46.9 | 54.5 | 551.7 | 86.5 |
| | $N(\mu_L + b_0, \sigma_L + b_0)$ | 13487.0 | 34.0 | 56.1 | 641.0 | 88.5 |
| FCD Var Intrahem | SHASH($\mu_L + b_0, \sigma_L + b_0, \nu, \tau$) | 13937.5 | 34.4 | 61.9 | 0.0 | 0.0 |
| | SHASH($\mu_S + b_0 + b_1x, \sigma + b_0, \nu, \tau$) | 13918.2 | 48.9 | 62.0 | 19.3 | 88.3 |
| | **SHASH**($\mu_L + b_0, \sigma + b_0, \nu, \tau$) | 13905.2 | 28.3 | 62.2 | 32.3 | 86.7 |
| | SHASH($\mu_S + b_0, \sigma + b_0, \nu, \tau$) | 13840.3 | 51.3 | 63.6 | 97.2 | 88.6 |
| | SHASH($\mu_L + b_0 + b_1x$) | -6274.1 | 0.0 | 48.7 | 20211.6 | 78.4 |

**Table S2.** Percentage of extreme deviations (*|Z| >* 1.645) across healthy test controls and clinical cases for static and dynamic functional connectivity (denoted by FC and FCD, respectively). Values are presented as *n/N* (%), where *n* is the number of extreme deviations and *N* is the total group size. To evaluate baseline calibration, local control groups were tested against the theoretical 10% expectation using exact Binomial tests (* *p_FDR_ <* 0.05). To evaluate clinical sensitivity, diagnostic groups were compared to their local, within-site control counterparts using Fisher’s Exact Tests. After correcting for multiple comparisons, no diagnostic group exhibited a statistically significant inflation in extreme deviations relative to local controls (all *p_FDR_ >* 0.05).

| Cohort | FC Inter-hem (%) |  | FC Intra-hem (%) |  | FCD Inter-hem (%) |  | FCD Intra-hem (%) |  |
| --- | --- | --- | --- | --- | --- | --- | --- | --- |
|  | Control | Case | Control | Case | Control | Case | Control | Case |
| HCP Young Adult | 77/636 (12.1%) | — | 76/636 (11.9%) | — | 56/633 (8.8%) | — | 57/633 (9.0%) | — |
| HCP Aging (HCA) | 89/748 (11.9%) | — | 85/748 (11.4%) | — | 60/748 (8.0%) | — | 58/748 (7.8%) | — |
| 1000BRAINS | 36/279 (12.9%) | — | 37/279 (13.3%) | — | 43/279 (15.4%)* | — | 39/279 (14.0%) | — |
| CAMCAN | 16/196 (8.2%) | — | 16/196 (8.2%) | — | 15/196 (7.7%) | — | 14/196 (7.1%) | — |
| ADNI | 2/29 (6.9%) | AD: 3/55 (5.4%)<br>MCI: 10/91 (11%) | 2/29 (6.9%) | AD: 2/55 (3.6%)<br>MCI: 11/91 (12.1%) | 3/29 (10.3%) | AD: 6/61 (9.8%)<br>MCI: 13/91 (14.3%) | 5/29 (17.2%) | AD: 7/61 (11.5%)<br>MCI: 18/91 (19.8%) |
| ABIDE | 12/132 (9.0%) | 35/374 (9.6%) | 9/132 (6.8%) | 32/374 (8.5%) | 10/65 (15.4%) | 24/195 (12.3%) | 7/65 (10.8%) | 29/195 (14.9%) |
| ADHD-200 | 4/84 (4.8%) | 19/215 (8.8%) | 7/84 (8.3%) | 23/215 (10.7%) | 5/42 (11.9%) | 10/102 (9.8%) | 5/42 (11.9%) | 17/102 (16.7%) |

## Notes

### Competing Interest Statement

The authors have declared no competing interest.

